# Chromoanagenesis is a driver of structural variation in the smallest photosynthetic eukaryote

**DOI:** 10.1101/2025.09.17.676765

**Authors:** Claire Bugnot, Tyler Alioto, Fernando Cruz Rodriguez, Jessica Gomez Garrido, Marta Gut, Sheree Yau, Gwenael Piganeau

## Abstract

Marine microalgae populations can rapidly evolve resistance to viruses upon infection. In *Ostreococcus mediterraneus* resistance to the prasinovirus OmV2 emerged within five days in all virus-exposed populations. Whole-genome sequencing of pairs of resistant and susceptible cell lines revealed extensive structural genomic changes, particularly on the Small Outlier Chromosome (SOC). SOC alterations included large deletions, duplications, rearrangements, and whole chromosome duplication, yet no consistent structural variant or single nucleotide polymorphism could be directly associated with resistance. Hybrid *de novo* assemblies confirmed the unique SOC assembly of each strain, with a highly polymorphic ∼2 kb tandem repeat region exhibiting an "accordion-like" pattern of expansion and contraction. No new viral insertions were found, though endogenous viral elements were conserved across lines. Two interchromosomal translocations between the SOC and chromosomes 2 and 17 provide evidence for chromoplexy, thereby offering novel insights into the mechanisms underlying the distinctive evolutionary path of this chromosome. Together, these findings demonstrate that resistance to OmV2 evolves rapidly and consistently but cannot yet be linked to any specific structural variations; instead, the high rate of localized genomic structural variations points to a distinct mechanism of chromosome evolution.

**Significance:** The mechanisms and consequences of structural variation remain poorly understood in many lineages, particularly in ecologically important marine microbes. This study shows that resistance to prasinovirus OmV2 in a cosmopolitan phytoplankton species, *Ostreococcus mediterraneus* evolves rapidly and consistently, even though no resistance-associated SNPs or shared structural variants were identified on standard chromosomes. Instead, extensive genomic rearrangements—particularly of a single chromosome—occur independently of viral infection. These findings highlight the role of chromoplexy as the driving mechanism underlying the diversification of the enigmatic Small Outlier Chromosome in this lineage.

## Introduction

Genetic diversity underlies environmental adaptation(Pellestor and Gatinois 2020). It arises both from micro-evolutionary processes—such as single nucleotide polymorphisms and small insertion-deletions (indels)—and macro-evolutionary processes, also known as structural variations (Alkan, Coe and Eichler 2011). Structural variations are typically defined as genomic differences larger than 50 bp, encompassing insertions, deletions, duplications, inversions, translocations and large tandem repeats (Alkan, Coe and Eichler 2011; Kosugi *et al*. 2023; Hu *et al*. 2025). Structural variations are induced by various molecular processes such as replication, repair mechanisms and DNA recombination (Currall *et al*. 2013). In some cases, these events can result in extensively rearranged chromosomes. Initially identified as the underlying cause of numerous genetic diseases in humans (Stankiewicz and Lupski 2010; Lupski 2023), structural variations are now also recognized as an important source of genetic diversity (Stankiewicz and Lupski 2010; Kloosterman and Cuppen 2013; Yi and Ju 2018; Koltsova *et al*. 2019; Peng *et al*. 2019; Dobry *et al*. 2023; Bian *et al*. 2024; Nair *et al*. 2025).

The decreasing cost and increasing accessibility of long-read sequencing technologies have enabled the characterisation of major chromosome rearrangements. Catastrophic chromosomal rearrangements have been collectively referred to as chromoanagenesis, from Greek ‘chromo’ meaning chromosome and ‘anagenesis’ meaning rebirth(Holland and Cleveland 2012). Three types of structural rearrangements have been identified from cancer cells: chromothripsis (‘thripsis’ for shattering(Stephens *et al*. 2011)), chromoanasynthesis (‘anagenesis’ for reconstruction(Liu *et al*. 2011)) and chromoplexy (‘pleko’ for to braid or to weave(Baca *et al*. 2013)). Chromothripsis corresponds to the shattering of a chromosome, or a chromosomal region, into tens to hundreds of fragments, which are later reassembled in a random order and orientation, generally involving non-homologous end joining (Stephens *et al*. 2011; Koltsova *et al*. 2019; Brás, Sebastião Rodrigues and Rueff 2020; Dewhurst 2020; de Groot *et al*. 2023). Chromoanasynthesis occurs on a single chromosome due to error-prone DNA replication, leading to copy number variations (Liu *et al*. 2011; Ostapińska, Styka and Lejman 2022; Guo, Comai and Henry 2023a). In contrast to the two aforementioned processes, chromoplexy involves multiple chromosomes with intra- and inter-chromosomal translocations (Baca *et al*. 2013; Guo, Comai and Henry 2023a).

Several intrinsic and extrinsic factors have been implicated in the genesis of these catastrophic genomic events (Holland and Cleveland 2012; Koltsova *et al*. 2019; Pellestor and Gatinois 2020; Guo, Comai and Henry 2023a). Endogenous mechanisms include cellular processes, such as micronucleation (Zhang *et al*. 2015), abortive apoptosis (Tubio and Estivill 2011), defects in the p53 pathway (Haferlach *et al*. 2008) and telomere crisis (Maciejowski *et al*. 2015). Additionally, environmental factors, such as exposure to ionizing radiation (Morishita *et al*. 2020), viral infections and genomic integration (Schütze *et al*. 2016; Koltsova *et al*. 2019), as well as artificial genome editing techniques such as CRISPR-Cas9 (Leibowitz *et al*. 2021) or biolistic transformation (Liu *et al*. 2019), have also been demonstrated to contribute to these events.

Initially discovered in human cancer cells, chromoanagenesis has since been documented across diverse eukaryotes, including mammals (Villagómez *et al*. 2008; Pinton *et al*. 2009; Carbone *et al*. 2014; Guo, Comai and Henry 2023a) and plants (Guo, Comai and Henry 2021, 2023b, 2023a). While it is often deleterious in animals, such genome restructuring seems to be more tolerated in plants, where it may contribute to adaptation, gene innovation, and genome evolution(Pellestor and Gatinois 2020). Similar large-scale genomic alterations have also been reported in model organisms such as *Saccharomyces cerevisiae* (Anand *et al*. 2014), *Caenorhabditis elegans* (Itani *et al*. 2016), as well as in the marine phytoplankton genus *Ostreococcus* (Blanc-Mathieu *et al*. 2017; Yau *et al*. 2020).

*Ostreococcus* is a genus of unicellular eukaryotic picoalgae (cell size < 2 µm) from the order Mamiellales (Chlorophyta) with a cosmopolitan distribution in the sunlit ocean (Delmont *et al*. 2022; Yung *et al*. 2022). Its compact haploid 13 Mb genome makes it a valuable model for studying chromoanagenesis-like events. It contains eighteen standard chromosomes and two so-called outlier chromosomes: the Big Outlier Chromosome (BOC) and the Small Outlier Chromosome (SOC) (Moreau *et al*. 2012). These chromosomes exhibit atypical features—low GC content, higher transposable element density, abundant repetitive sequences, and reduced gene density— relative to the rest of the genome (Moreau *et al*. 2012). While the BOC is the mating type determination candidate (Blanc-Mathieu *et al*. 2017), the SOC is thought to play a role in antiviral defense. The hypervariability of the SOC in natural populations, along with its mosaic structure, suggests that *Ostreococcus* undergoes—or tolerates—chromosomal reshuffling events reminiscent of chromoanagenesis(Blanc-Mathieu *et al*. 2017). This underscores its potential as a model for investigating the origin and consequences of chromosome instability in unicellular eukaryotes.

*Ostreococcus* is infected by prasinoviruses, which are double-stranded DNA viruses classified within the phylum *Nucleocytoviricota*, also known as nucleocytoplasmic large DNA viruses (NCLDVs), which are extremely abundant in their natural environment (Meng *et al*. 2021). These viruses replicate within the host, assembling in the cytoplasm and ultimately causing cell lysis (Dunigan, Fitzgerald and Van Etten 2006; Derelle *et al*. 2008; Weynberg, Allen and Wilson 2017). Interestingly, *Ostreoccocus* populations develop reversible resistance to prasinoviruses, allowing for long-term coexistence(Yau *et al*. 2020). A population is considered susceptible (S) if a significant proportion of its cells can be infected, leading to lysis and viral production. However, after lysis, the population recovers systematically (Yau *et al*. 2016). Once a recovered population no longer undergoes lysis upon re-exposure to the virus, it is considered resistant (R). Resistant populations primarily consist of uninfected R cells, although viral production still occurs at low levels, probably due to infection of a minority of susceptible cells forming a coexistence dynamic, akin to a bet-hedging strategy between susceptible and resistant cells (Yau *et al*. 2020). The molecular mechanisms underlying the emergence of anti-viral resistance remain poorly understood. Previous work demonstrated the systematic evolution of resistance to viruses in *O. tauri* and its association with karyotypic changes in the SOC (Yau *et al*. 2016, 2018). Intraspecific genome comparison revealed that the length of the SOC was positively correlated with resistance to a broader spectrum of prasinovirus strains (Blanc-Mathieu *et al*. 2017). Reciprocally, the transition from a resistant parental line to susceptible experimental evolution lines has been associated with large deletions on the SOC in the species *O. mediterraneus* (Yau *et al*. 2020).

This study aimed to elucidate the genetic variation associated with the evolution of resistance to viruses in *O. mediterraneus* by identifying structural variations, as well as point mutations, linked to susceptible or resistant phenotypes, and the presence or absence of viral sequence insertion. Genomic analyses of five independent pairs of virus-susceptible control and evolved-resistant lines of *O. mediterraneus* revealed substantial structural variation within the SOC. Surprisingly, no evidence was found for an association between this variation and the resistant phenotype, while sequence analyses highlighted a high rate of spontaneous structural rearrangements relative to point mutations in this chromosome.

## Results

### *Systematic evolution of resistance to prasinovirus OmV2 in* Ostreococcus mediterraneus

To understand the genomic basis of resistance acquisition to prasinovirus, an experimental evolution study was conducted with 40 independent culture lines, derived from a single cell of the virus-susceptible *O. mediterraneus* strain MA24 (Yau *et al*. 2020). Of these, 32 lines were split into infected and uninfected control flasks, while 8 lines remained as non-infected controls (Fig. 1A). All infected cultures visibly lysed within 3 days post-infection, but resistance to prasinovirus systematically evolved in all 32 infected lines, as shown by regrowth within five days post-infection. Whole genome sequences of five selected pairs of resistant (R) and susceptible control (S) lines, designated lines A to E, were obtained using Illumina short-read and Oxford Nanopore Technology long-read sequencing.

**Fig. 1:**
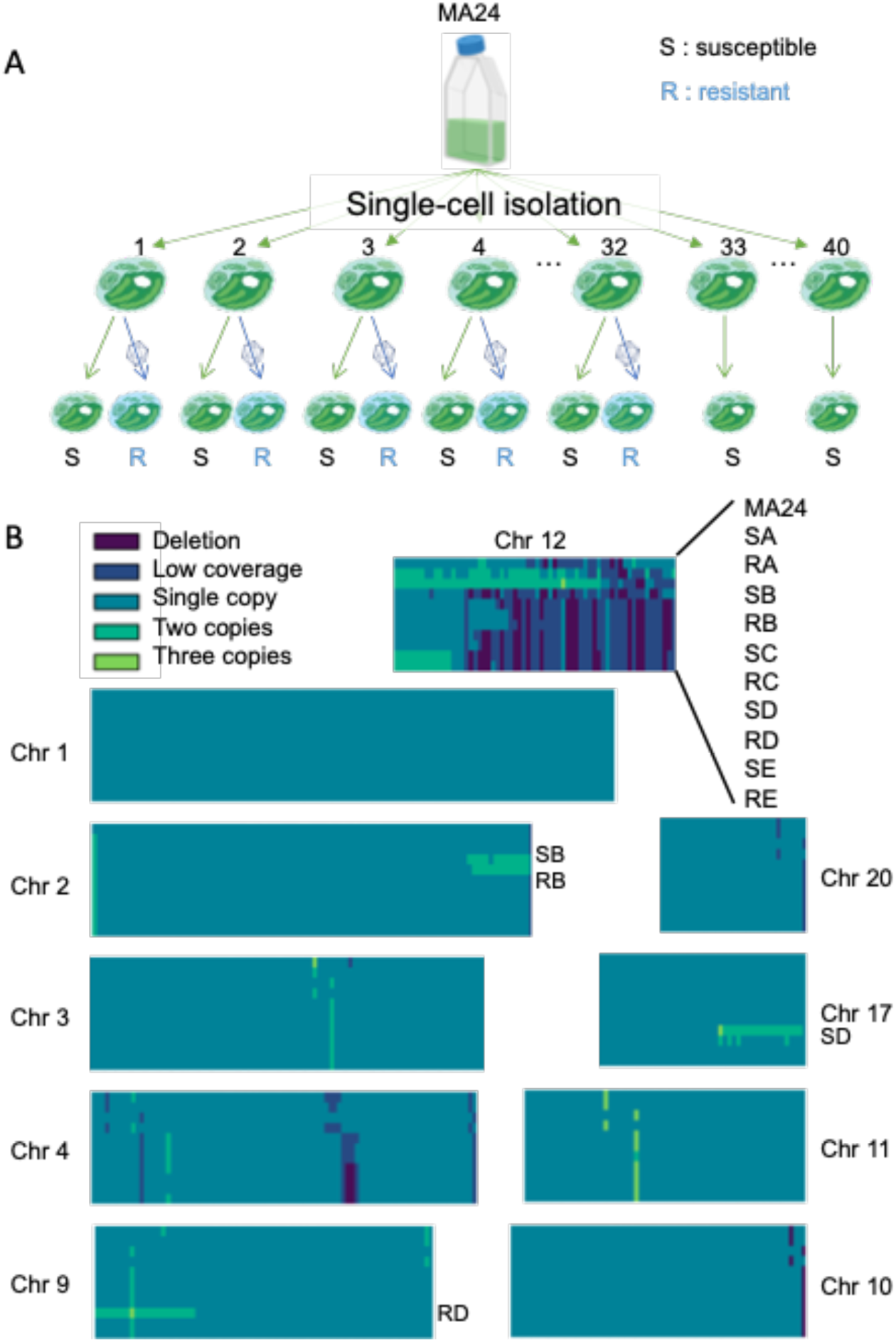
Detection of copy number variations in the genome of *O. mediterraneus* from virus-susceptible and resistant experimental-evolution lines. **A:** Illustration of the experimental evolution study setup. Single-cells from the parent virus-susceptible line (MA24) were sorted into 40 culture lines, 32 of which were divided into pairs to which the prasinovirus OmV2 was added to one of each pair. All lines challenged with OmV2 lysed, regrew and developed resistance. **B:** Genome-wide copy number variations in the parent MA24 and the five experimental-evolution lines analysed, comprising virus-susceptible (S) and virus-resistant (R) pairs with each line designated as A–E. The data from each experimental line were stacked for each chromosome. Copy number variations were estimated by short read mapping onto the reference *O. mediterraneus* RCC2590 genome, with the read coverage of each chromosome normalized to the coverage of chromosome 5. Coverage was estimated over 10 kb windows. The 10 chromosomes with the most copy number variations and chromosome 1 with no variations are displayed (the complete heatmaps are available in Fig. S1). Specific lines with large copy number variations in chromosomes 2, 9 and 17 are labelled.

### Line specific and shared copy number variations on standard chromosomes

The experimental evolution lines (A–E) and the parental line (MA24) were screened to identify copy number variations relative to the ancestral reference line RCC2590 by analyzing chromosome coverage in 10 kb windows using the Illumina short-reads. Most of the 20 chromosomes showed no large copy number variations (Fig. S1). The only copy number variation consistently shared across all five R and S line pairs was a 10 kb duplication at the left end of chromosome 2 (Fig. 1B). Beyond this shared event, seven standard chromosomes exhibited notable copy number variations, most prominently chromosomes 2, 9, and 17 (Fig. 1B). On chromosome 2, both B line strains (SB and RB) carried a 120 kb duplication at the right end of the chromosome. Notably, the D lines displayed two large copy number variations greater than 100 kb: strain RD had a 240 kb duplication at the end of chromosome 9, including a 10 kb triplication, while strain SD carried a 170 kb duplication at the end of chromosome 17. Apart from the aforementioned copy number variations, the R and S strains in lines C, D and E exhibited similar copy number variation patterns across the genome. Likewise, SA and SB showed shared copy number variations and overlapped significantly with those found in the parental line MA24. However, the most substantial copy number variations—both deletions and duplications—across all strains were concentrated on a single chromosome: chromosome 12, also known as the Small Outlier Chromosome (SOC) in this species.

### The SOC as a hotspot of line-specific copy number variations

Most copy number variations between strains were concentrated on the SOC. A striking observation was the presence of large low coverage blocks spanning ∼400 kb region at one end of the SOC, indicating extensive deletions concentrated in this region across all lines. We compared the size and positions of these deletions in the parental line MA24 and the ten experimental evolution lines relative to the 637 kb SOC of the reference strain RCC2590. MA24 exhibited a 220 kb deletion between positions 300-520 kb. Interestingly, portions of this deleted region in MA24 were retained in all experimental evolution lines, particularly in line A. This pattern suggests a deletion polymorphism in the parental population, which was later differentially assorted among the experimental evolution lines. Supporting this, pulsed field gel electrophoresis from Yau *et al*., 2020(Yau *et al*. 2020) showed that MA24 harbored at least two SOC variants, one approximately 453 kb and another around 200 kb in size, consistent with the presence of two SOC deletion variants in the parental MA24 population. Thus, strain RA, which had the smallest deletion (∼150 kb, Fig. 2), likely derived from a lower frequent SOC variant in the MA24 population. In contrast, the largest deletion (∼480 kb) was found in the E strains, suggesting an additional SOC deletion event occurred after the divergence from the parental line.

**Fig. 2:**
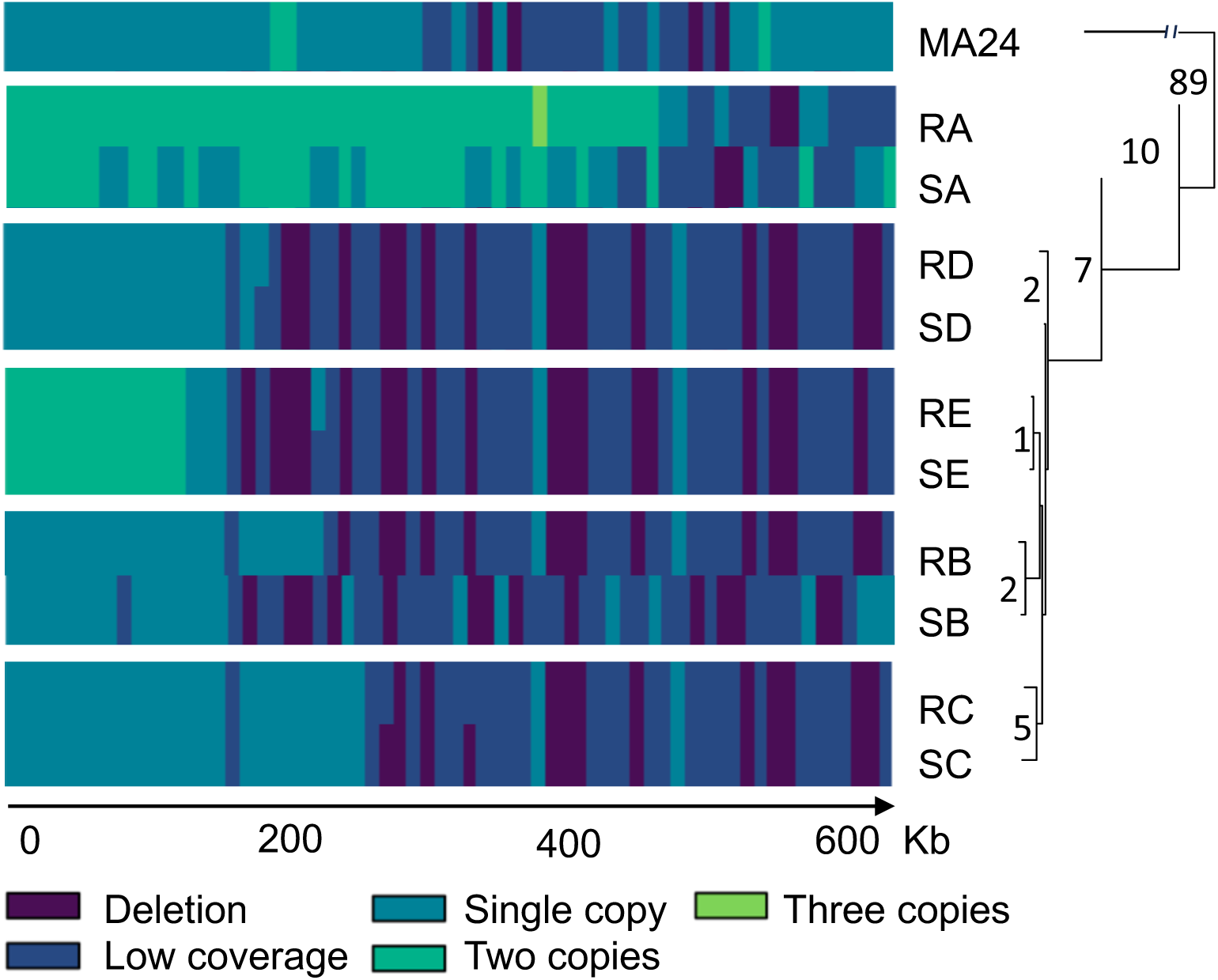
Copy number variation of the SOC compared with clustering analysis based on the distance matrix between strains based on single nucleotide polymorphisms (SNPs) in the MA24 parental line and the five pairs of experimental evolution lines (A–E). Left: Zoom-in on the coverage of the SOC from Fig. 1 showing deletions and duplications relative to the 653 kb reference genome. Right: dendrogram showing the clustering of the 11 strains based on 120 SNPs detected in the callable genome, excluding the SOC. The exact number of SNPs between each pair of lines is indicated on the phylogenetic tree.

The second striking observation was a duplication at the left end of the SOC in strains A and E. In the A strains, nearly the entire 470 kb region at that end was duplicated, with a segment between positions 380–390 kb in RA suggesting a triplication. In the E strains, a 130 kb region at the same end was duplicated. Additionally, SOC coverage patterns revealed that independent copy number variations occurred between the resistant and susceptible members of each A–E line pair, suggesting these rearrangements likely arose after the evolution of resistance to prasinovirus OmV2. Despite this, SOC coverage profiles indicated that strains primarily clustered by the experimental lineage map (Fig. 1A), and no SOC variant could be consistently associated with either the resistant or susceptible phenotype.

### Single Nucleotide Variation reflects the experimental lineage map, not resistance to OmV2

To investigate whether any single nucleotide polymorphisms (SNPs) were associated with the R or S phenotypes, we identified SNPs by mapping of Illumina short-reads from each line to the RCC2590 reference genome. A total of 120 SNPs were located within the callable genome, defined as regions covered in all 10 strains and thus excluding the highly repetitive SOC (see Methods). The number of SNPs per line ranged from 75 in line RA to 91 in line SE (Table S1), with 19 SNPs detected in the parental line MA24. Within pairs, the number of SNP differences varied from just one SNP in the SE/RE pair to as many as 10 between SA and RA. A phylogenetic tree based on SNP distances largely reflected the experimental lineage map, with S and R pairs clustering together rather than by phenotype, except for strain RA, which diverged from this pattern (Fig. 2). Importantly, no SNPs were found to be uniquely associated with either the R or the S phenotype on the callable genome.

In conclusion, neither SOC copy number variants nor SNP distributions could be directly associated with the development of resistance to prasinovirus OmV2. Instead, we observed large insertions and deletions suggestive of spontaneous structural variations of the SOC, occurring independently of viral infection and acquisition of resistance.

### Hybrid genome assemblies resolve structural variants predicted by copy number variation

To more precisely characterize these structural variations, all SOCs were reconstructed using hybrid *de novo* assemblies combining long- and short-read sequencing, yielding complete or near-complete chromosomal assemblies (Fig. 3). Each SOC assembly was validated by remapping both short and long reads to confirm sequence coverage and scaffold placement (Table 1). The SOCs of the 10 experimental evolution lines grouped into four distinct structural types.

**Fig. 3:**
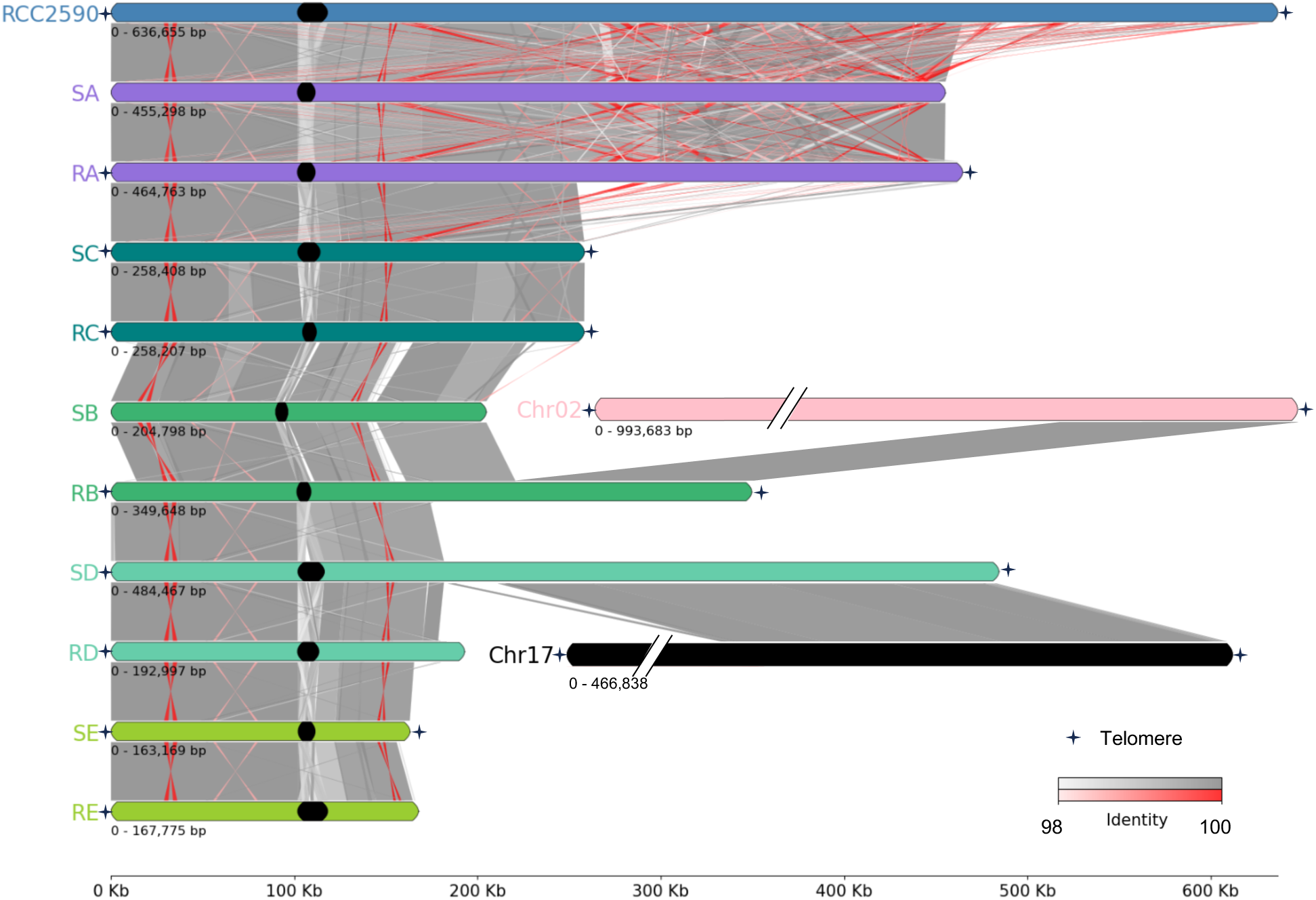
Comparison of the SOC assemblies of the experimental evolution lines and the reference RCC2590. High nucleotide identity (98%–100%) regions between pairs of chromosome assemblies are joined with shaded grey (same orientation) and red (reverse orientation) blocks. SOC assemblies from the same experimental evolution line pair (R/S) share the same color. Inter-chromosomal translocations are shown by alignment of the SOC to the reference chromosome 2 (Chr02) and chromosome 17 (Chr17). The black lozenge on the SOC assemblies shows the size and location of a repeat region.

**Table 1.** Assembly features of each small outlier chromosome.

| Strain line | Name of contigs | RCC2590<br>order | Telomeres | Bridging<br>long reads |
| --- | --- | --- | --- | --- |
| SA | Reverse<br>AP0970.masurca_flye2.5_s24 | 1 | 0 | 158 |
|  | AP0970.masurca_flye2.5_s22 | 2 | 0 |  |
| SB | Reverse<br>AP0971.masurca_flye2.5_s098 | 1 | 0 | 20 |
|  | AP0971.masurca_flye2.5_s063 | 2 | 0 | 19 |
|  | Reverse<br>AP0971.masurca_flye2.5_s106 | 3 | 0 |  |
|  | Reverse<br>AP0971.masurca_flye2.5_s093 | 4 | 0 | 10 |
|  | AP0971.masurca_flye2.5_s072 | 5 | 0 | 4 |
| SC | AP0974.masurca_flye2.5_s25 | 1 | 1 | 42 |
|  | AP0974.masurca_flye2.5_s24 | 2 | 1 |  |
| SD | AP0967.masurca_flye2.5_s15 | 1 | 2 | 35 |
| SE | AP0976.masurca_flye2.5_s31 | 1 | 1 | 124 |
|  | AP0976.masurca_flye2.5_s34 | 2 | 1 |  |
| RA | AP0969_SOC | 1 | 2 | - |
| RB | AP0972.masurca_flye2.5_s13 | 1 | 2 | 19 |
| RC | Reverse<br>AP0973.masurca_flye2.5_s50 | 1 | 1 | 26 |
|  | AP0973.masurca_flye2.5_s60 | 2 | 0 | 28 |
|  | AP0973.masurca_flye2.5_s52 | 3 | 0 |  |
|  | Reverse<br>AP0973.masurca_flye2.5_s76 | 4 | 0 | 27 |
|  | Reverse<br>AP0973.masurca_flye2.5_s65 | 5 | 1 | 20 |
| RD | Reverse<br>AP0968.masurca_flye2.5_s29 | 1 | 1 | 29 |
|  | Reverse<br>AP0968.masurca_flye2.5_s33 | 2 | 0 | 16 |
|  | Reverse<br>AP0968.masurca_flye2.5_s42 | 3 | 0 |  |
| RE | AP0975.masurca_flye2.5_s25 | 1 | 1 | 18 |
|  | Reverse<br>AP0975.masurca_flye2.5_s33 | 2 | 0 |  |

The first type of assembly spanned the entire chromosome from telomere-to-telomere and included the SC and RC lines, as well as RA and SE. These SOCs exhibited a large deletion at one end compared to the reference, consistent with previous coverage analyses. Their sizes were 258 kb, 258.5 kb, 426 kb and 163 kb, respectively, much shorter than the length of the 637 kb reference SOC.

The second type comprised SOCs with a single telomere (RD and RE). The SOCs of RD and RE (193 kb and 168 kb long, respectively) contained a telomere at the left end and exhibited large deletions of 444 and 470 kb at the right end, also consistent with the coverage analyses.

The third assembly type corresponded to the reconstructed SOC of line SA and SB, which lacked identifiable telomeric repeats. SB has a size of 205 kb, presenting a deletion of 15 kb on the left end and one of 417 kb at the right end. For SA, despite the absence of telomeres, the left end was identical to those of other SOC assemblies, and the right end exhibited a similar large deletion similar to its paired line, RA. This was consistent with coverage analyses, suggesting that the SA SOC assembly was nearly complete. The final estimated size was 455 kb, with a 145 kb deletion at the right end.

The fourth type of assembly included the SOCs of strains RB and SD, both of which formed telomere-to-telomere assemblies that, unexpectedly, incorporated sequences from another chromosome. In both cases, the left end of the chromosome was from the SOC. In strain RB, the assembly joined 220 kb of the SOC with a duplicated and translocated 120 kb segment from the right end of chromosome 2. Coverage analyses confirmed this duplication on chromosome 2 (Fig. 1B). In strain SD, 181 kb of the SOC’s left end was joined to 300 kb from the left end of chromosome 17. For both *de novo* assemblies, visual inspection of long and short read mappings confirmed the junctions between the SOC and the respective translocated chromosomal segments. Their paired lines SB and RD, appeared to exhibit only a deletion at the 3′ end of the SOC (Fig. 3). Interestingly, in both RB and SD, the SOC assemblies revealed that intrachromosomal translocations account for the copy number variation of part of chromosome 2 in RB, and part of chromosome 17 in SD.

There is, however, one discrepancy between the coverage analyses and the *de novo* assemblies concerning the 5′ duplications observed in lines A and E: the size of the duplicated part identified in the coverage analyses closely matches the size of the respective SOC assemblies. As no other chromosomes in these lines showed evidence of duplication, we inferred that lines A and E each harbor two copies of the SOC.

### Conservation of OmV2 insertions across experimental evolution lines

Given recent evidence of endogenous viral elements in algal genomes (Moniruzzaman *et al*. 2020; Erazo-Garcia *et al*. 2025), the genome assemblies of the reference and the 10 evolution experimental lines were screened for insertion of DNA from the OmV2 virus. In the reference genome (RCC2590), five OmV2-derived insertions were identified, located on chromosome 4 (73 bp), chromosome 6 (58 bp), chromosome 8 (80 bp), chromosome 10 (36 bp) and chromosome 11 (339 bp). These five insertions were conserved across all experimental lines, except for the chromosome 4 OmV2-derived insertion in SB. No new OmV2 insertions were identified in any of the 10 experimental evolution lines.

### Accordion-like dynamics of polymorphic tandem repeats

One region of the SOC assemblies, consisting of 3 to 8 tandem repeats of an approximately 1.8 kb unit (Fig. 4) presented exceptional dynamics. The minimal structural rearrangement of this region observed within a pair of lines was observed in RA and SA with an identical number of 5 repeats. A similar region had previously been identified in the SOC of a natural *O. tauri* population (Yau *et al*. 2016; Blanc-Mathieu *et al*. 2017). However, there was no significant local alignment between the *O. tauri* and *O. mediterraneu*s repeats. The repeats occurred in tandem (Fig. 4), with polymorphisms among copies. Repeat sizes ranged from 1,685 to 1,985 bp, with size variation primarily due to small terminal deletions on the final repeat, with the terminal repeat in line RD truncated to 777 bp.

**Fig. 4:**
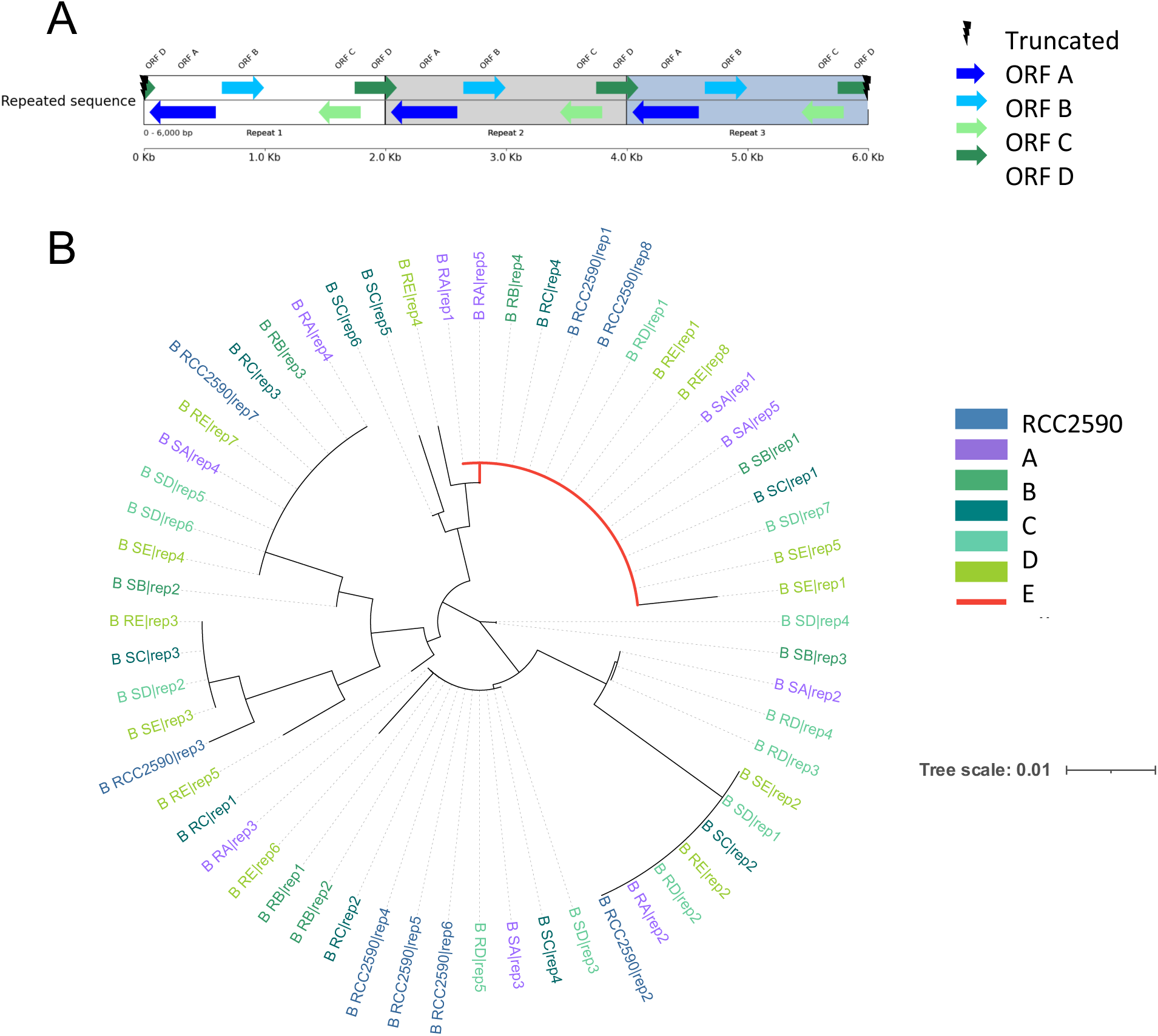
Features of the tandem repeat ‘accordion’ region and inference of its evolutionary history in the experimental evolution and reference lines. A: Schematic representation of three copies of the accordion region showing the relative positions of predicted ORFs. B: Phylogenetic tree using neighbour-joining of the 114 amino acids of all the copies of ORF B in the 11 lines (experimental evolution lines A–E and reference RCC2590). The red branch line highlights the only identical copy present in all strains.

Some of the repeats were interrupted by a 1,001 bp spacer sequence, increasing the total size of the composite repeat unit to 2,929 bp. This spacer contained long inverted repeats of 225 bp but did not encode any identifiable open reading frames (ORFs). The same spacer sequence was detected six times in the reference genome—on chromosomes 4, 7, 8, 9, 10, and within the SOC. Within the polymorphic tandem repeat regions, the spacer was present only once per array. Although the spacer appeared in the consensus sequences of lines SA, SB, SC, RD, and RE, long-read analysis revealed its presence in all SOC assemblies except that of line SD. The position of the spacer-containing repeat varied, with one to three possible insertion sites across the population, accounting for its absence in some consensus sequences. Based on its consistent presence and polymorphic position, this spacer may represent a non-autonomous transposable element that inserted prior to the expansion of the tandem repeat arrays. Its inactivity is suggested by the constant single copy of the spacer, despite variation in repeat number through duplication or deletion.

During this study, the various sequence similarity searches (see method) of the entire polymorphic tandem repeated regions, or the repeats themselves, did not allow the identification of any similar sequences in GenBank (nr, nr/nt), suggesting these repeats are fast evolving and specific to *O. mediterraneus*.

Candidate ORFs were detected within all the polymorphic tandem repeated regions. None of these ORFs contained known functional domains, as determined by InterProScan analysis. The ORFs were designated A, B, C, and D according to their relative positions within the repeat unit (Fig. 4A). An additional reverse ORF, named E, was identified in the SD line, located between the ORFs B and C. Each of the four primary ORF groups included both a more conserved form and variants characterized by point mutations and indels.

Further protein structure predictions suggested that ORFs C and the E may contain a transmembrane domain. For the remaining ORFs, structural predictions were inconclusive. Notably, a gene annotated in the *O. mediterraneus* genome (Ostme30g0062), composed of two exons and one intron, corresponds in both sequence and genomic position to a combination of ORF C and E. The predicted structure of Ostme30g0062 matches the structure of a predicted monomeric transmembrane protein of *Micromonas commoda* and of a predicted monomer forming the trimeric transmembrane protein in *Ostreococcus lucimarinus* and *Micromonas pusilla*. This structure conservation suggests that the sequence may play a functional role shared among related marine algae.

Since ORF B was the only ORF consistently present in all but the truncated terminal repeat, it was used to investigate the evolutionary history of the tandem repeats within and between the strains. An alignment of the 114 amino acids of the ORF B copies from all strains was used to construct a phylogenetic tree (Fig. 4B). The resulting phylogeny revealed that copies from all the strains had mixed origins, with some likely inherited vertically and others having diversified independently within specific lineages. Only one ORF B copy was identical across all strains (highlighted in red in Fig. 4B); however, this copy did not occupy the same position in the repeat array in each strain, indicating polymorphic rearrangements in repeat order among strains. Phylogenetic trees constructed from alignments of the most conserved versions of ORFs A, C, and D supported this pattern of strain-specific expansion, contraction, and rearrangement of the repeat structures.

To assess whether the predicted ORFs in the repeat units were likely to encode functional proteins under selection on amino-acid composition, we estimated the number of synonymous and non-synonymous mutations among ORF copies (Table S2). These analyses did not allow us to reject the null hypothesis of an equal rate of synonymous and non-synonymous changes, consistent with a lack of selection on the amino-acid sequences of these putative ORFs. Therefore, ORFs A, B, C and D are unlikely to encode protein coding sequences.

Overall, this region was shown to be highly dynamic, with lineage-specific repeats with modifications suggesting a high point mutation rate, as well as cycles of expansion and contraction—an accordion-like mechanism shaping repeat copy number and order.

## Discussion

### Structural variations point towards unique chromosome instability in one chromosome

We resolved large structural variations and point mutations in the genomes of both susceptible and resistant experimental evolution lines. The dynamics of chromosome 12, the SOC, stood out, as it exhibited the most extreme variation in read coverage among strains. Large-scale deletions at its 3′ end were observed across all lines, confirming previous observations based on pulsed-field gel electrophoresis and hybridization (Yau *et al*. 2020). The ten outlier chromosome assemblies revealed large deletions, duplications, including whole chromosome duplications, intra- and inter-chromosomal translocations, resulting in unique SOC assemblies for each of the 10 lines.

The SOC had been previously characterized in all complete genome sequences of Mamiellales sequenced to date (Derelle *et al*. 2006; Worden *et al*. 2009; Moreau *et al*. 2012), whose species divergence is estimated to date back more than 300 million years (Yung *et al*. 2022), and aneuploidies involving the SOC have been previously reported in mutation accumulation lines of *Micromonas* and *Bathycoccus* (Krasovec *et al*. 2023). Additionally, population genomic analyses in one species revealed a striking contrast: whereas standard chromosomes show approximately 1% nucleotide divergence between any pair of individuals, only 15–30% of the SOC chromosome could be aligned between individuals from the same population (Blanc-Mathieu *et al*. 2017).

While this novel type of hypervariability was proposed to result from a chromothripsis-like process (Blanc-Mathieu *et al*. 2017), the structural rearrangements we describe here are not consistent with chromothripsis as the evolutionary mechanism for this chromosome. Instead, the process resembles partial chromosome duplication and chromoplexy, involving only a few catastrophic events rather than the chromosome shattering characteristic of chromothripsis (Guo, Comai and Henry 2023a).

### Large chromosome duplications and chromoplexy drives diversification of the SOC

The high rate of duplication, deletion, and interchromosomal fusion observed on this chromosome within the approximately 190 generations separating each pair of lines, thus helps us better understand the evolutionary dynamics of this outlier chromosome.

First, chromoplexy explains the high number of strain-specific genes on this chromosome, i.e. genes uniquely present in one strain and not in other strains from the same species (Blanc-Mathieu *et al*. 2017). Indeed, frequent random fusion with parts of different chromosomes, followed by deletions, causes rapid diversification of the gene content between strains on this chromosome within a population. Using this new information about the origin of the SOC, we specifically searched for sequence homologies between the standard chromosomes and the SOC, while excluding repeated elements that occur in multiple genomic locations. This approach led to the identification of 25 signatures of past translocation events, comprising a total of 77 Kb within the SOC that share significant sequence identity with regions of the standard chromosomes. Notably, this includes a 36 Kb segment originating from chromosome 4. On the longer timescale of speciation, chromoplexy is thus expected to gradually lead to the loss of synteny on this chromosome (Fig. 5), which has been reported in early comparative genome studies in *Ostreococcus* and *Bathycoccus* (Palenik *et al*. 2007; Moreau *et al*. 2012).

**Fig. 5:**
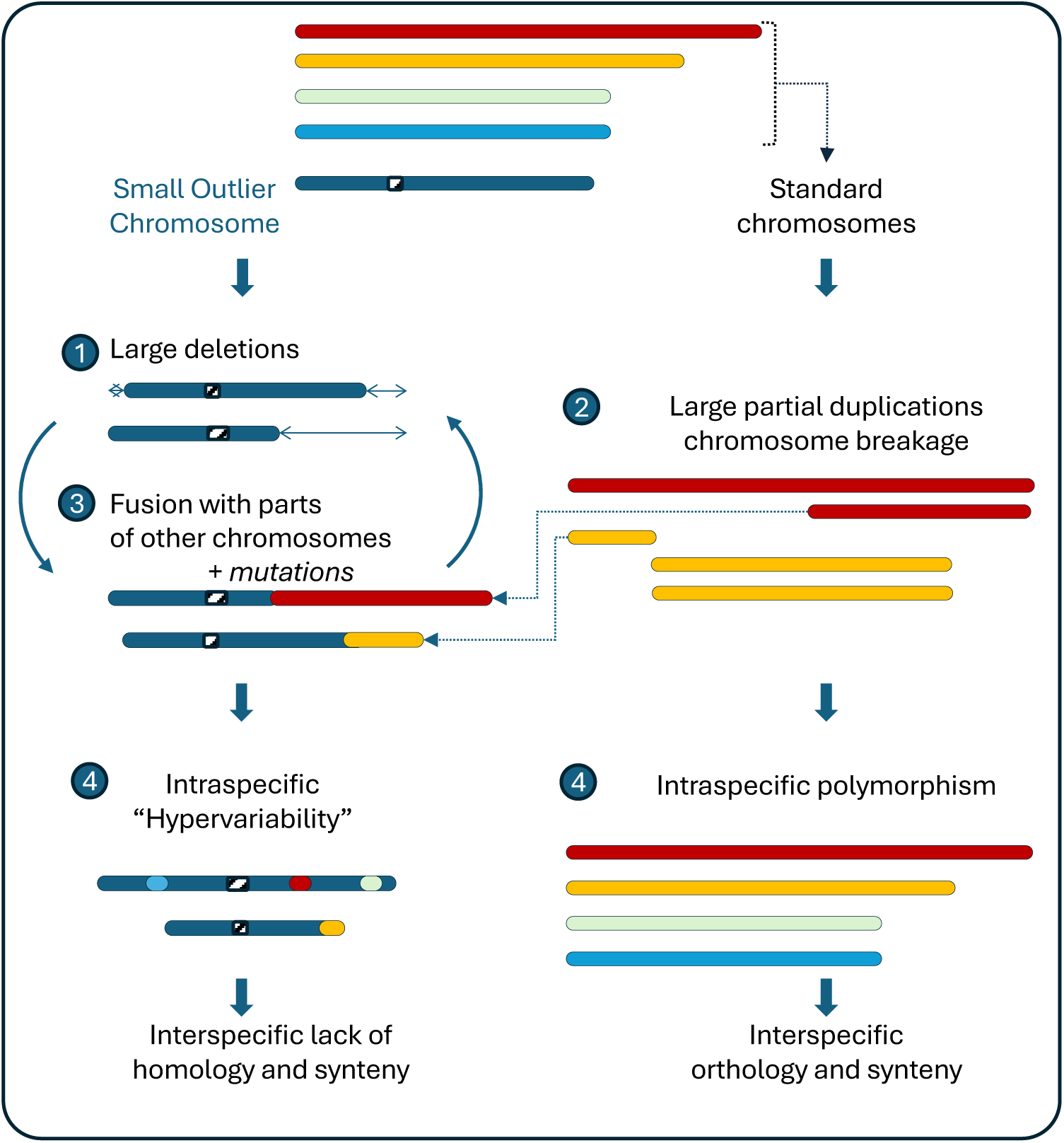
Model of hypervariable chromosome evolution driven by large deletions, fusions with parts of duplicated chromosomes, and mutations (left part), as opposed to the evolution of standard chromosomes (right part).

More precisely, the present experiment provides insights into the relative tempo of structural variation on this chromosome versus point mutations on the standard chromosomes: during the course of these five independent infection and resistance evolution experiments, the ten strains have accumulated between one and ten independent point mutations on standard chromosomes between pairs (Fig. 2). These can be compared to the total number of independent structural mutations on the SOC. Structural variations between each pair of strains corresponded to a minimum of—counting the minimum number of indels and accordion repeat copy number variations—ten duplications-deletions larger than 10 kb and two large chromosomal translocations from chromosome 17 (300 kb) and 2 (120 kb). Therefore, the number of large structural variations (12) on this chromosome is of the same order of magnitude as the number of point mutations on all 18 standard chromosomes (20). The spontaneous mutation rate in the smallest photosynthetic eukaryote corresponds to approximately one point mutation every 100 cell divisions (Krasovec *et al*. 2017). This, in turn, allows us to calibrate the structural divergence observed between two strains with the number of nucleotides differences we observe between two strains. Under the hypothesis that the structural variations of the SOC observed in *O. mediterraneus* can be extended to other Mamiellales species, the 50,000 point mutations between two pairs of strains in a natural population of *O. tauri* (Blanc-Mathieu *et al*. 2017), would correspond to approximately the same number of structural changes on the SOC. This is consistent with the low level of synteny on the SOC among closely related strains. Because of the high divergence time between each Mamiellales species sequenced so far and the saturation of substitutions at synonymous positions, equivalent to more than one substitution at each neutral site, this relative timeline also accounts for the complete lack of synteny on this chromosome observed between species (Palenik *et al*. 2007; Worden *et al*. 2009; Moreau *et al*. 2012).

### Sequence divergence in genes located on the SOC

Another feature of the SOC is the sharp decrease in the proportion of orthologous genes located on this chromosome (Jancek *et al*. 2008). This pattern could be explained by three, non-mutually exclusive, scenarios: (i) a high turnover of gene content driven by the integration of sequences from other lineages, such as viruses; (ii) an elevated point mutation rate obscuring sequence identity; or (iii) an elevated fixation rate of mutations as a consequence of reduced recombination. These three possibilities are discussed in detail below.

Indeed, green algae frequently show evidence of viral sequence insertions, including from giant viruses (NCLDVs) and virophages (Sheng *et al*. 2022), with some integrations being quite extensive (Moniruzzaman *et al*. 2020). However, no newly inserted OmV2 sequences were detected during this experiment. This argues against the hypothesis of the SOC being a hotspot for ongoing viral integration—at least in *O. mediterraneus*. It suggests instead that prasinoviruses either rarely integrate into this genome, or that viral sequences are efficiently eliminated following genome integration.

Nevertheless, viral-derived sequences are not entirely absent. Previous studies identified viral segments in both the standard and SOC of *O. tauri* (Blanc-Mathieu *et al*. 2017), and we also detected short prasinovirus-like sequences within the standard chromosomes of *O. mediterraneus*. While these fragments are limited, their presence suggests ancient or low-frequency integration events, even if the SOC is not an active hotspot of viral integration in this species.

Thus, an elevated point mutation rate on genes on the SOC seems the more likely explanation for the low rate of orthologous relationships on genes located on this chromosome. Estimating the point mutation rate is notoriously difficult in highly repetitive regions, which are typically excluded from spontaneous mutations estimates (Krasovec *et al*. 2017, 2023). However, our analysis of the polymorphism of the accordion region suggests many more point mutations occur within this region than within the standard chromosomes, and this would be consistent with an elevated nucleotide mutation rate on this chromosome. Another plausible evolutionary scenario over a longer timescale is that the lack of recombination in such rearranged regions promotes the evolution of genomic islands of divergence, a pattern that has been reported in many species (Sodeland *et al*. 2016; Dobry *et al*. 2023; Marín-García *et al*. 2024; Akopyan *et al*. 2025).

### Are chromosome copy number variation, chromoplexy and transcriptional reprogramming of the SOC connected?

In the nucleus of eukaryotic cells, chromosomes are folded into subnuclear domains known as chromosome territories (Cremer *et al*. 2006; Meldi and Brickner 2011). This folding is dynamic, allowing for the formation of chromatin loops that facilitate both inter- and intra-chromosomal interactions involved in gene coregulation (Simonis *et al*. 2006; Meldi and Brickner 2011). The spatial positioning of chromatin within the nucleus influences transcriptional activity: repressed regions are often localized at the nuclear periphery, while reactivation is associated with their relocation to the nucleoplasm (Ragoczy *et al*. 2003; Reddy *et al*. 2008; Meldi and Brickner 2011). The SOC has a bipartite transcriptional organization (Yau *et al*. 2016), with the accordion region demarcating the two parts of the chromosomes being transcriptionally active or inactive in different strains. The SOC may therefore have up to 80% of its chromatin in a transcriptionally repressed state in some strains (Yau *et al*. 2016). Therefore, a large part of the non-transcribed SOC is expected to be located within the nuclear periphery and membrane.

This, in turn, may contribute to a higher rate of structural variation, as impaired migration during mitosis can lead to micronucleus formation. Chromosomes trapped within micronuclei are exposed to a more damaging environment, making them more susceptible to chromoanagenesis (Leibowitz, Zhang and Pellman 2015; Koltsova *et al*. 2019; Guo, Comai and Henry 2023a). Additionally, the formation of chromatin loops extending beyond chromosome territories for interchromosomal gene coregulation may facilitate interchromosomal rearrangements. To date, micronuclei have never been reported in these species. However, recent chromosome Hi-C analyses indeed suggest an insulated position for the SOC in *O. tauri*(Valiadi *et al*. 2025). Therefore, a deeper understanding of nuclear organization is poised to provide valuable insights into genome plasticity, particularly concerning the unique characteristics of the SOC.

### A highly polymorphic accordion region lacking conclusive evidence of protein-coding potential

While the structural rearrangements underlying both intraspecific hypervariability and interspecific synteny loss have now been characterized, the role of this chromosome in conferring resistance to prasinoviruses remains unclear. Our results demonstrate that chromosomal rearrangements occur independently of viral infection. Although we extended previous observations of systematic resistance evolution following viral infection in O. *tauri* (Yau *et al*. 2016) to *O. mediterraneus*, we did not identify any common genome rearrangement associated with the five resistant strains. The only systematic changes between S and R were in the sequence of the SOC and the accordion region, either in the number of tandem repeats or in mutations affecting the number of predicted open reading frames (ORFs).

The core of the accordion region consists of a long polymorphic sequence repeated in tandem, with copy numbers ranging from 3 to 8. However, the number of copies does not appear to correlate with the resistance phenotype: one lineage (A) had identical copy numbers in S and R strains, two lineages (B and E) had more copies in R strains, while two others (C and D) had more in S strains. If gene copy number in this region is associated with the phenotype, it is unclear whether these genes encode for protein coding genes or non-coding RNAs. Among the open reading frames identified in the accordion region, ORF B was consistently found across all strains and all repeat copies, yet it shares no homology to any known protein-coding gene.

Interestingly, the only open reading frame in the accordion region with a predicted protein structure is a fusion of ORFs C and E, which likely encodes a transmembrane protein (Ostme30g0062). As homologous genes can be identified in three other species, this open reading frame likely encodes a protein sequence.

The lack of evidence for other protein coding genes for the ORF in this region points towards the encoding of functional alternatives such as non-coding RNAs. Non-coding RNA longer than 200 nucleotides (LncRNAs) that do not code for proteins, typically evolve rapidly, show low sequence conservation, and are often expressed in a highly context-specific manner. Their evolutionary dynamics are strongly influenced by genomic features such as bidirectional promoters and transposable elements, both of which are known to facilitate the gain and loss of lncRNA loci (Kapusta and Feschotte 2014). As such, genomes rich in transposable elements tend to produce more flexible and rapidly evolving transcriptomes. In the nucleus, lncRNAs can modulate chromatin structure by interacting with chromatin-remodeling complexes, either by guiding, scaffolding, or sequestering them to specific genomic loci (Han and Chang 2015; Neve *et al*. 2020). LncRNAs are thus good candidates to be encoded within the transcriptionally dynamic SOC, and especially in the transcriptomic rewiring around the accordion region, which corresponds to the physical limit between repressed and expressed regions of the SOC (Yau *et al*. 2016).

In conclusion, we resolved the structural dynamics of the SOC, characterized by spontaneous extensive tandem repeat polymorphisms and structural rearrangements, including translocations from other chromosomes. The inherent instability of the SOC highlights its potential as a key player in fast evolution, raising intriguing questions about the molecular mechanisms involved and the broader implications for the origin and resolution of spontaneous complex chromosomal rearrangements in eukaryotes. Future studies focusing on transcriptomic and proteomic analyses, as well as chromatin spatial organization, are poised to shed more light on the molecular mechanisms underlying the relationship between transcription, structural variation, and viral resistance in *Ostreococcus mediterraneus*.

## Materials and methods

### Algal strains, culture conditions and experimental evolution experiment

In this study, all the strains belong to the species *O. mediterraneus*. The parent line of all strains, “MA24”, (S3 line in Yau *et al*., 2020(Yau *et al*. 2020)) was obtained by mutation accumulation on the reference strain RCC2590 (Krasovec *et al*. 2017). Strains were maintained in culture in L1 medium prepared using autoclaved seawater, filtered at 0.22 µm. They were kept under 12:12 hour light/dark regime using white light at 15°C. Forty cultures of MA24 were grown from a single cell of the MA24 strain following the protocol described in Krasovec *et al*., 2016(Krasovec *et al*. 2016, 2017). Thirty-two cultures were divided into two flasks and one flask of each strain was infected with the prasinovirus OmV2, following the infection procedure previously described in Yau *et al*., 2020 (Yau *et al*. 2020). After cell lysis, resistance to OmV2 developed in all infected cultures within 21 days. Eighteen resistant cultures survived until sequencing. Thus, 18 pairs of susceptible and resistant-evolved cultures were available, with independent evolution over 84 days since the viral infection (9/09/2019 to 2/12/2019). They were referred to as S and R in the manuscript. A PFGE was performed on 24 samples corresponding to 12 (R, S) pairs, following the protocol described previously(Yau *et al*. 2020). The analysis of this PFGE led us to select five (R, S) pairs for Illumina and Nanopore sequencing. The study is focused on 5 pairs of R and S culture lines that had developed resistance to the viruses, namely lines 6, 10, 14, 26, 28, renamed A, B, C, D and E, respectively.

### DNA extraction and sequencing

DNA extraction and sequencing were performed as described in Thomy *et al*., 2021(Thomy *et al*. 2021). Briefly, DNA was extracted from five strains (A to E) for both phenotypes using the CTAB (cetyltrimethylammonium bromide) protocol. DNA quality was assessed using the absorbance ratios at 260/280 nm and 260/230 nm (Nanodrop), visualized by gel electrophoresis on a 0.8% agarose gel, and quantified by fluorometry (Quantus). The library was prepared using a modified KAPA HyperPrep kit protocol, then quality-checked and quantified. Illumina and Oxford Nanopore Technologies sequencing were conducted on the 10 samples as described in Thomy *et al*., 2021 (Thomy *et al*. 2021).

### Genome assemblies and Small Outlier Chromosome (SOC) reconstruction

Raw Illumina paired-end libraries and Oxford Nanopore Technologies reads were used with MaSuRCa v3.4.1 (Zimin *et al*. 2013) and Flye v2.5 (Kolmogorov *et al*. 2019), which respectively construct mega-reads and assemble them, to obtain hybrid assembly.

To improve the assembly of the SOC, contigs belonging to the SOC (completely or partially) were identified using nucmer (mummer-4.0.0beta2), mummerplot and show-coords (Marçais *et al*. 2018). The identified contigs were used to reconstruct the chromosome with the RCC2590 SOC as reference (Geneious Prime 2023.0.4, https://www.geneious.com). Without overlap, the contigs were placed using the reference for orientation, thus ensuring a conservative estimation of the structural variation within the data. The reconstructions were validated through visual inspection in IGV (Robinson et al. 2011) of both long read and short read alignments, resulting in high confidence assemblies. The features of the SOC reconstruction are summarized in Table 1. The presence of virus sequences in the SOC’s assemblies was checked by BLASTing the OmV2 genome against the assemblies. To visually check the structural rearrangements and avoid false positive structural variants (David *et al*. 2024), the reconstructed SOC were aligned against each other with blastn (length_thr=600, identity_thr=98) and the identity between strains was represented using PyGenomeViz (Shimoyama 2024) within a range of 98-100%. Lower identity was considered as non-conserved sequences.

### Structural Variations: read coverage analyses

The short reads were mapped against the RCC2590 reference genome using bwa-mem2 (version 2.2, mem) (Md *et al*. 2019). The coverage for windows of 10 kb was obtained with samtools (samtools-1.9, view, sort and index) and bedtools (bedtools2-2.18.0, makewindows and coverage). The coverage dataset was used to create a heatmap (Rstudio 2022.07.2, https://www.R-project.org, heatmap) against the reference genome RCC2590 to obtain a visual representation of coverage variations, after normalizing coverage by the read coverage of a standard invariant chromosome (chr 5). Windows where fewer than 10% of positions were covered by at least one read were categorized as deletions. Windows with single, two-fold, or three-fold coverage were delineated by merging all normalised coverage values across all strains and chromosomes. The 90% confidence interval for single-copy coverage was estimated using the qt function in RStudio (Rstudio 2022.07.2), resulting in a range of 56% to 140% of the mean coverage. The same interval was applied to define two-fold (140% to 224%) and three-fold overage (224% to 308%.) windows. Windows with more than 10% but less than 56% of covered positions were categorized as low coverage (Fig. 1, 2, S1).

### SNPs analyses on the callable genome

Raw Illumina reads were quality checked with FastQC (FastQC_v0.11.7) and compared with MultiQC (MultiQC-1.14) (Andrews 2010; Ewels *et al*. 2016). The reads were trimmed with fastp (fastp-0.23.4) to remove adapters (--adapter_fasta), N base (-n 0) and to conserve only reads longer than 80bp (-l 80) (Chen *et al*. 2018; Chen 2023). The trimmed reads were mapped again, against RCC2590 with bwa-mem2 (version 2.2, mem) (Md *et al*. 2019). The bam format was obtained using samtools (samtools-1.9, view, sort and index). Then, the duplicates were flagged using gatk MarkDuplicates (version 4.4.0.0) (Depristo *et al*. 2011). The SNPs were called with bcftools (version 1.17) mpileup, call (--ploidy 1) and filter (‘QUAL>20 && DP>100’). The vcf files were indexed with tabix. The callable genome was considered to be all the chromosomes except the SOC because of its repeat content. Since the sequencing came from a population of cells rather than a single cell, only SNPs with at 90% frequency were kept. The DP4 parameter and the localization of the SNPs in the vcf files were used to select the SNPs above this frequency threshold. A matrix of all the 120 SNPs of the 11 samples identified was compiled. A binary matrix was created by generating a binary result (0: absence of the SNPs, 1: presence of the SNPs) by checking all the samples and the parent line’s SNPs against the SNPs matrix. This matrix was then treated with R studio (Rstudio 2022.07.2) using the Kimura (1980) distance (hclust) to obtain the SNP phylogenetic dendrogram.

### Analysis of the ’accordion’ region of the SOC

The identification of a pattern was done using a dotplot of the RCC2590 reference SOC sequence against itself (Geneious Prime 2023.0.4). The analysis was extended to all the stains SOC and the *O. tauri* RCC4221 SOC and confirmed by the long read alignment. A blast of the RCC2590 pattern sequence against the RCC4221 pattern sequence was done to identify homologous sequences between the two species patterns. The main component of the pattern was identified as a long polymorphic sequence repeated in tandem using Geneious Prime 2023.0.4. The repeated sequence and the whole polymorphic tandem repeated region were blasted (blastn, tblastx, blastx) against the NCBI database (nt, nr/nt, internal transcripted spacer region) to identify homologous sequences. ORFinder (https://www.ncbi.nlm.nih.gov/orffinder/) was used to determine candidate ORFs in the whole polymorphic tandem repeated region. Candidates ORF were classified depending on the position in the repeated sequence and the candidate amino-acid sequence. Each type of candidate ORF observed in all strains was aligned (global alignment with free end gaps, cost matrix of 93% similarity, gap open penalty 12, gap extension penalty 3) and respective phylogenetic trees were built (Jukes-Cantor distance using neighbor-joining, iTOL 7.0 (Letunic and Bork 2024), as implemented in Geneious Prime 2023.0.4. The position and copy number of the spacer sequence were validated by long read mapping against the assemblies. The protein structure of each type of candidates ORF was determined by alphafold 3 (Abramson *et al*. 2024) and was compared to known or predicted structures using Foldseek (van Kempen *et al*. 2023). Domain and function were check with InterProScan (Blum *et al*. 2025).

## Resource availability

The Genbank accession number of the *O. mediterraneus* reference genome is GCA_012295225.1, and the accession number of OmV2 MN688676.

The accession numbers of the data generated and analyzed in this study are in project PRJEB9049, the ONT data accession numbers are from ERR15283078 to ERR1528387, and the Illumina data accession numbers: from ERR15283067 to ERR15283076.

Requests for further information and resources should be directed to and will be fulfilled by the lead contact, Gwenael Piganeau.

## Acknowlegments

We would like to thank all GENOPHY Lab members, in particular Amandin James, Marc Krasovec, Lisa Mettrop, and Nigel Grimsley, for their stimulating discussions and support. We also thank Frederic Sanchez and Adrien Cadoudal for their help with the experiments and DNA extractions. We acknowledge the GenoToul Bioinformatics platform (Toulouse, France) for bioinformatic analysis support and cluster availability. CNAG acknowledges the support of the Spanish Ministry of Science and Innovation through the Instituto de Salud Carlos III and the 2014–2020 Smart Growth Operating Program and cofinancing with the European Regional Development Fund (MINECO/FEDER, BIO2015-71792-P). We also acknowledge the support of the Generalitat de Catalunya through the Departament de Salut and Departament d’Empresa i Coneixement.

## Declaration of AI used

During the preparation of this work the authors used ChatGTP to correct English grammatical errors. After using this tool, the authors reviewed and edited the content as needed and take full responsibility for the content of the publication.

